# Type III interferons disrupt the lung epithelial barrier upon viral recognition

**DOI:** 10.1101/2020.05.05.077867

**Authors:** Achille Broggi, Sreya Ghosh, Benedetta Sposito, Roberto Spreafico, Fabio Balzarini, Antonino Lo Cascio, Nicola Clementi, Maria De Santis, Nicasio Mancini, Francesca Granucci, Ivan Zanoni

## Abstract

Lower respiratory tract infections are a leading cause of mortality driven by infectious agents. RNA viruses such as influenza virus, respiratory syncytial virus and the new pandemic coronavirus SARS-CoV-2 can be highly pathogenic. Clinical and experimental evidence indicate that most severe and lethal cases do not depend on the viral burden and are, instead, characterized by an aberrant immune response. In this work we assessed how the innate immune response contributes to the pathogenesis of RNA virus infections. We demonstrate that type III interferons produced by dendritic cells in the lung in response to viral recognition cause barrier damage and compromise the host tissue tolerance. In particular, type III interferons inhibit tissue repair and lung epithelial cell proliferation, causing susceptibility to lethal bacterial superinfections. Overall, our data give a strong mandate to rethink the pathophysiological roles of this group of interferons and their possible use in the clinical practice against endemic as well as emerging viral infections.

---

The ability to resolve viral infections of the lung is dependent on the actions of interferons (IFNs) and inflammatory cytokines. These molecules are induced during viral infections, yet the relative contributions of each interferon or cytokine to host defense and a return to homeostasis remains undefined. In particular the actions of type III IFNs (IFN-λ) have attracted much attention, as these cytokines operate primarily at mucosal surfaces, in part due to the selective expression of the type III IFN receptor in epithelia and neutrophils (*1–3*). Recent work established that unlike their type I or II counterparts, type III IFN signaling induces antiviral activities while simultaneously minimizing the tissue destructive activity of neutrophils (*4–6*). When considered in the context of an increasing number of viral infections, including the recently emerged severe acute respiratory syndrome (SARS)-coronavirus (CoV)-2, where inflammation appears to be the primary driver of life threatening symptoms (*7–12*), the ability of type III IFNs to limit immunopathology but maintain antiviral activity is significant. Indeed, discussions on the possible use of IFN-λ against SARS-COV-2 have begun (*13*) and a clinical trial (*14*) has been initiated with the human pegylated-IFN-λ1a. Despite this interest of the use of type III IFNs to treat viral infections, we know very little about the short- and long-term actions of this family of secreted factors.

During viral infections of the lung, immunopathology may predispose the host to opportunistic bacterial infections, referred to as superinfections, which take advantage of this unbalanced status and often lead to lethal consequences (*15–17*). In contrast to its protective effect against acute viral infections, IFN-λ was found to decrease bacterial control in virus-bacteria superinfection models(*18–20*). Whether this is due to the anti-inflammatory activity of IFN-λ that results in a reduced resistance of the host to a subsequent bacterial infection or to the capacity of this group of interferons to alter the physiology of the lung upon a viral encounter remains unresolved. As of present the contribution of type III IFN to the human pathology of severe viral infections and bacterial superinfections is unknown. Superinfections represent the first cause of lethality upon influenza virus infection (*21*) and correlate with severity in coronavirus disease (COVID)-19 patients (*11, 22–24*). Mouse models of SARS, middle east respiratory syndrome (MERS) (*25, 26*) and influenza (*5, 27–29*) are characterized by a robust induction of IFN-λ, however, the involvement of this group of interferons in COVID-19 is more controversial. Although *in vitro* studies on nasal and bronchial primary epithelial cells demonstrate IFN-λ induction by SARS-CoV-2 (*30*), another study found a reduced production of both type I and type III IFNs *in vitro* and *ex vivo* (*31*).

To directly evaluate the capacity of SARS-CoV-2 to induce interferons in the upper or lower airways, we tested the presence of IFNs, and other inflammatory cytokines, in the swabs of COVID-19 patients and healthy controls, as well as the bronchoalveolar (BAL) fluid of severe SARS-CoV-2-positive patients. Although we confirmed that COVID-19 patients present a scarce production of inflammatory mediators in the upper airways, the BAL fluid of patients with severe disease presented elevated levels not only of inflammatory cytokines, but also of specific members of the type III IFN family (Fig. 1A-E). Thus, IFN-λ production is associated with the most pathogenic endemic as well as emerging RNA viral infections.

**Figure 1.**
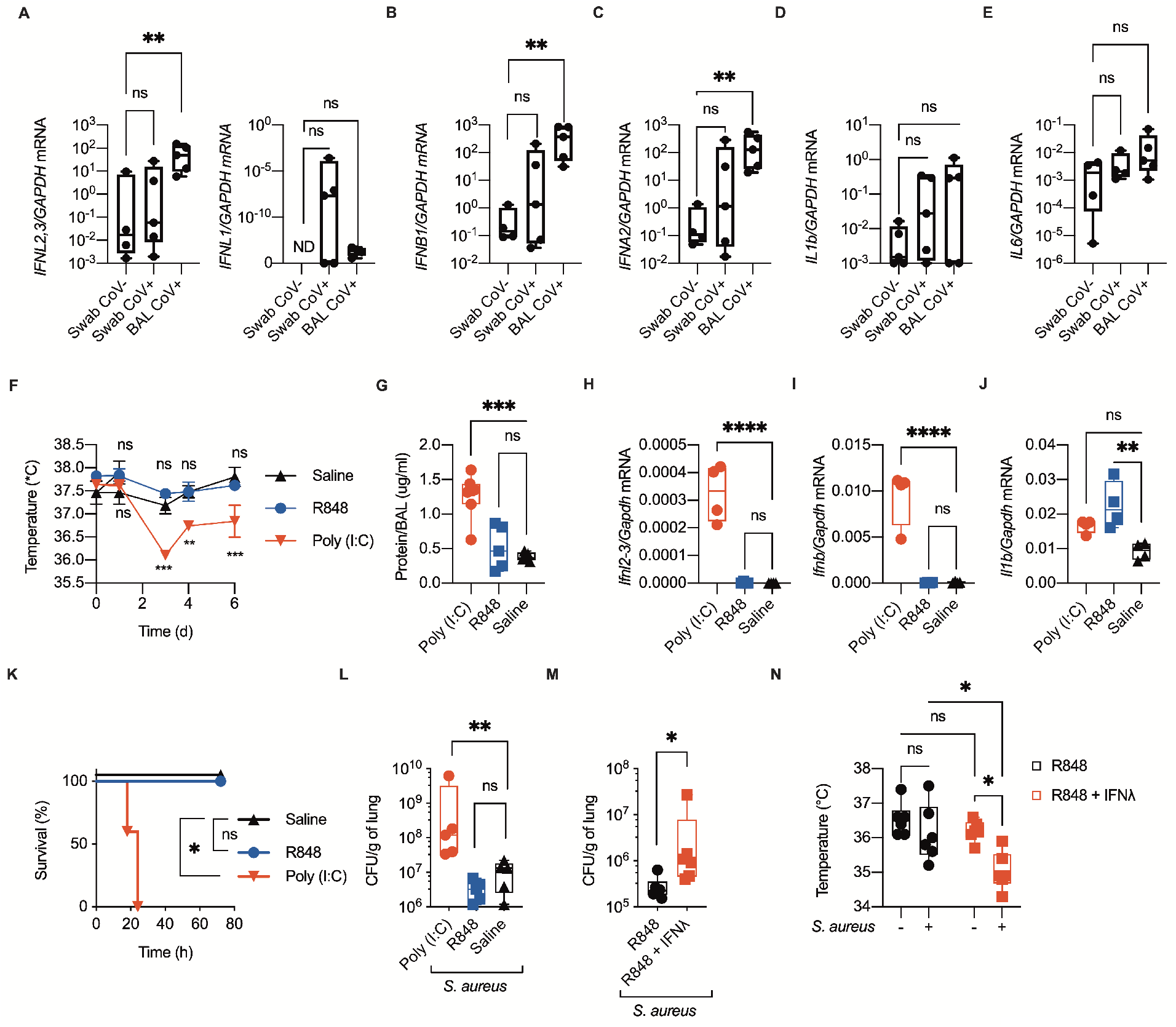
Morbidity correlates with the high expression of IFN-I and IFN-λ in the lung of COVID-19 patients’ BAL and of poly (I:C)-treated mice. (A-E) mRNA expression of *IFNL2,3, IFNL1* (A), *IFNB* (B), *IFNA2* (C), *IL1B* (D), and *IL6* (E) were evaluated in naso-oropharyngeal swabs from SARS-CoV-2-positive (Swab CoV+) and -negative (Swab CoV-) subjects and from bronchoalveolar lavages of intensive care unit (ICU)-hospitalized SARS-CoV-2-positive patients (BAL CoV+). (F-J) Mice were treated daily with i.t. 0.5 mg/kg poly (I:C), 0.5 mg/kg R848 or saline for 6 days. (F) Body temperatures of the treated mice measured over time are represented. (G) Amount of total protein in the BAL was measured at day 6 post treatment. (H-J) *Ifnl2,3* (H), *Ifnb1* (I), *Il1b* (J) mRNA expression was assessed in total lung lysate harvested 6 days post treatment. (K, L) Mice treated as in (F-J) were infected at day 6 with 0.5 × 10^8^ CFU of *S. aureus* administered i.t. and were monitored for survival (K). Bacterial load in the lungs of the treated mice normalized to lung weight was assessed 12 hours post infection (hpi) (L). Mice were administered daily i.t. R848 (0.5 mg/kg) or a combination of R848 and IFN-λ (50μg/kg) for 6 days and were then infected as in (K). Bacterial burden in the lungs (M) and body temperatures (N) before and after *S. aureus* infection are depicted. (G-J, L-N) Each symbol represents one mouse, median and range are represented. (F) Mean and SEM of 5 mice per group is represented. (K) Survival plot of 5 mice per group. (F-N) Representative data of 3 independent experiments. Statistics: ns, not significant (P > 0.05); *P < 0.05, **P < 0.01 and ***P < 0.001. Two-way ANOVA (F, N), One-way ANOVA (G-J, L), or two-tailed T test (M) were performed. Logarithmic values were fitted when evaluating bacterial load (L, M). Log-rank (Mantel-Cox) test, corrected for multiple comparisons was performed to evaluate survival (K).

In order to directly evaluate the contribution of IFN-λ to the immune pathology driven by RNA respiratory viruses uncoupled from its effect on viral replication, we devised an experimental system in which pattern recognition receptors (PRR) involved in viral sensing were stimulated with their cognate ligands in mice that can, or cannot, respond to IFN-λ. RNA viruses are directly sensed by two major classes of PRR (*32, 33*). Toll-like receptors (TLR) recognize viral nucleic acids in the endosomal compartment with TLR7 recognizing single-stranded RNA directly from virions in the endosomes (*34*) and TLR3 recognizing double-stranded RNA intermediate derived from dying infected cells (*35*). The cytosolic sensors retinoic acid-inducible gene I (RIG-I) and melanoma differentiation-associated protein 5 (MDA5) instead, recognize double strand and hairpin RNA from replicating viruses (*32, 33*) in association with the common adaptor mitochondrial antiviral signaling protein (MAVS). We thus intra-tracheally instilled the TLR7 ligand R848 and the synthetic analog of double-stranded RNA, polyinosine-polycytidylic acid (poly (I:C)) that stimulates both TLR3 and the RIG-I/MDA5-MAVS pathway *in vivo* (*36, 37*). Viral-sensing PPRs were stimulated over the course of six days to simulate prolonged innate immune activation in the lung. Only poly (I:C) treatment induced morbidity as measured by body weight and temperature drop (Fig. 1F and Fig. S1A), and only poly (I:C) treatment compromised barrier function as measured by the level of total protein and lactate dehydrogenase (LDH) in the bronchoalveolar lavage (BAL) (Fig. 1G and Fig. S1B). Increase in morbidity and decrease in barrier function correlated with the levels of IFN-I and IFN-λ, whose mRNAs are strongly upregulated in the lung of mice treated with poly (I:C) but not in mice treated with R848 (Fig. 1H, I). In contrast R848 treatment induced upregulation of pro inflammatory cytokines (i.e. IL-1β) but had no correlation with barrier function or temperature decrease (Fig. 1F-J).

Alterations in the epithelial barrier are known to predispose to lethal bacterial superinfections (*38–40*), we therefore infected mice treated with either R848 or poly (I:C) with *Staphylococcus aureus (S. aureus*). Only poly (I:C) treatment was sufficient to induce lethality upon *S. aureus* infection (Fig. 1K). Moreover poly (I:C) treated mice displayed a higher bacterial burden (Fig. 1L), a stronger hypothermia upon *S. aureus* infection, and higher barrier damage (Fig. S2A, B). *S. aureus* infection did not alter the pattern of IFN production as only poly (I:C) induces upregulation of interferons’ mRNAs (Fig. S2C-E). While both IFN-I and IFN-λ transcripts are upregulated over time upon poly (I:C) treatment (Fig. S3A, B), IFN-I protein levels measured in whole lung lysates are low and plateau after 1d of treatment while IFN-λ protein levels increase over time (Fig. S3C, D) and their increase correlates with *S. aureus* bacterial burden (Fig. S3E). Moreover, transcript levels of CXCL10, which is induced by IFN-I and not by IFN-λ (*5, 41*) peak at day 1 and reach a plateau over the following days (Fig. S3F), consistent with IFN-I protein expression kinetics. Conversely, the expression of the interferon-stimulated gene (ISG) *Rsad2*, which is induced similarly by both IFN classes, increases over time and, thus, correlates with increasing IFN-λ production (Fig. S3G). As expected, pro-inflammatory cytokines are expressed at low levels throughout the kinetic study (Fig. S3H, I). The correlation of bacterial burden with IFN-λ protein kinetics suggests a causal relationship between IFN-λ production and susceptibility to superinfection. Indeed, administration of exogenous IFN-λ concomitantly with R848, which induces immune activation but not interferon production, was sufficient to induce sensitivity to *S. aureus* infection, increasing morbidity and bacterial burden (Fig. 1M, N). Overall, these data indicate that IFN-λ is sufficient to break the lung tolerance and induce susceptibility to subsequent bacterial infections.

Next, we examined if IFN-λ is not only sufficient but also necessary to establish morbidity and sensitize to bacterial superinfection upon viral exposure. Contrary to wild-type (WT) mice, mice deficient in interferon lambda receptor 1 expression (*Ifrnlr1*^−/−^ mice) are completely protected from poly (I:C) induced morbidity, both in terms of hypothermia induction and weight loss (Fig. 2A, and Fig. S4A), and do not develop barrier damage (Fig. 2B, and Fig. S4B). Consistently with the absence of barrier defects, *Ifnrl1*^−/−^ mice are completely protected from lethality upon *S. aureus* superinfection (Fig. 2C) and show lower bacterial burden (Fig. 2D), lower barrier damage and decreased hypothermic response (Fig. 2E, F) compared to WT mice. Deletion of IFNLR1 doesn’t impact levels of mRNA or protein of IFNs or pro-inflammatory cytokines (Fig. S4C-H), confirming the direct role of IFN-λ in this context. Collectively these observations show that sustained expression of IFN-λ after RNA virus recognition is both necessary and sufficient to induce barrier breach and sensitization to *S. aureus* infection.

**Figure 2.**
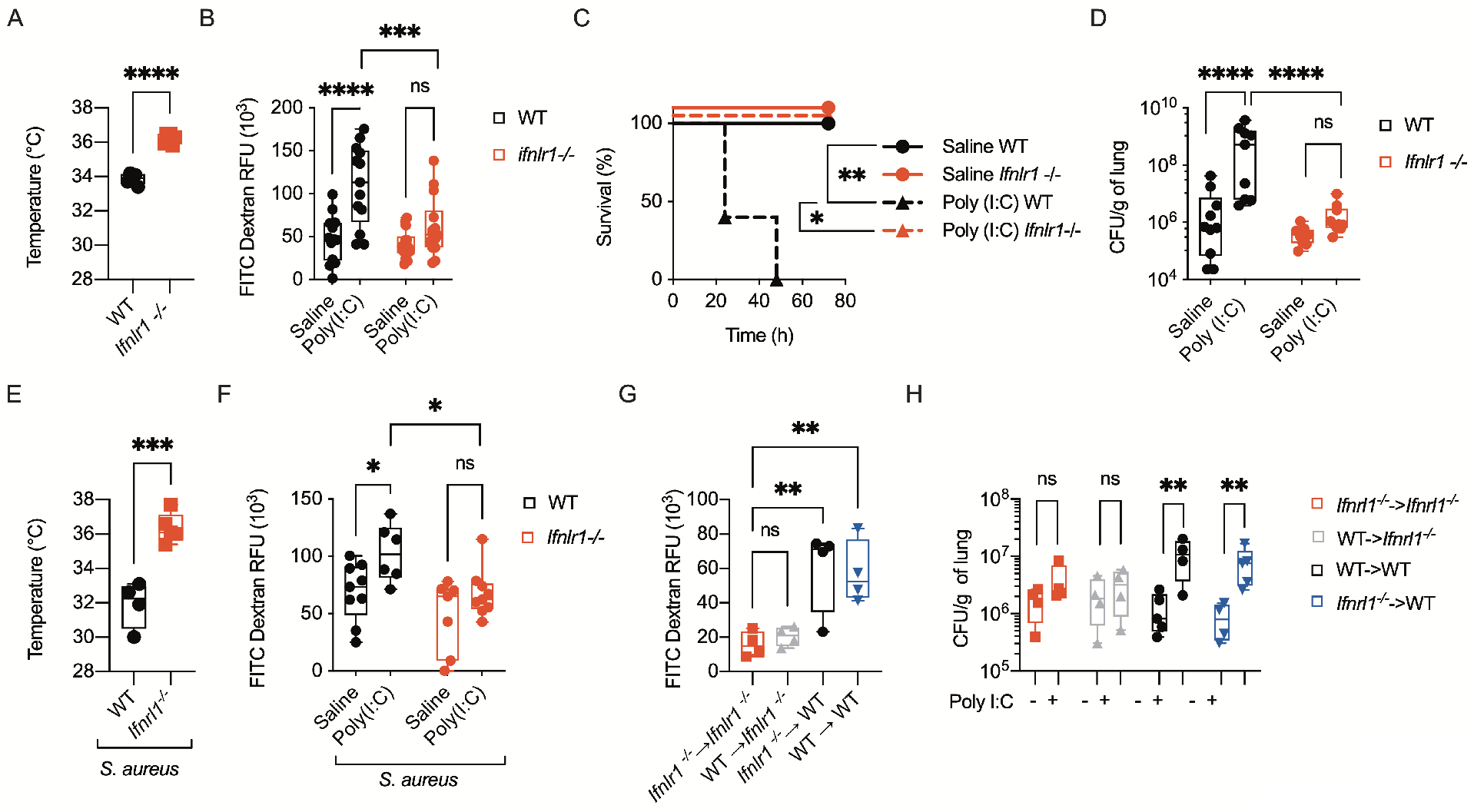
IFN-λ is necessary to increase susceptibility to bacterial infection induced by antiviral immunity. (A, B) WT and *Ifnlr1*^−/−^ mice were treated daily with 0.5 mg/kg poly (I:C) or saline for 6 days. (A) Body temperatures of poly (I:C)-treated WT and *Ifnlr1*^−/−^ mice were recorded on day 6. (B) On day 6 mice were treated i.t. with FITC-Dextran (10μg/mouse). Barrier permeability was measured as relative fluorescent units (RFU) of FITC-Dextran leaked in plasma 1 hour after injection. (C-F) WT and *Ifnlr1*^−/−^ mice treated with i.t. 0.5 mg/kg poly (I:C) or saline for 6 days, were infected i.t. with 0.5 × 10^8^ CFU of *S. aureus* and monitored for survival (C). Lung bacterial burden normalized by lung weight (D), body temperature (E), and barrier permeability (F) (as in (B)) were assessed 12 hpi. (G-H) Lethally irradiated WT or *Ifnlr1*^−/−^ recipients were reconstituted with donor bone marrow (*Ifnlr1*^−/−^ or WT) for 6 weeks and were treated as in (C-F). (G) Barrier permeability measured (as in (B)) and (H) lung bacterial burdens were evaluated 12 hpi. Each symbol represents one mouse, median and range are represented. (C) Survival plot of 5 mice per group. (A-H) Representative data of 3 independent experiments. Statistics: ns, not significant (P > 0.05); *P < 0.05, **P < 0.01 and ***P < 0.001. Two-way ANOVA (B, D, F, H)), One-way ANOVA (G), or two-tailed T test (A, E) were performed. Logarithmic values were fitted when evaluating bacterial load (D, H). Log-rank (Mantel-Cox) test, corrected for multiple comparisons was performed to evaluate survival (C).

Our data demonstrate that IFN-λ alters the lung epithelial barrier function, but we and others also showed that this group of interferons downmodulates host immune resistance, by inhibiting ROS release, migration, and pro-inflammatory cytokine release by neutrophils(*4–6*), a key immune cell population involved in the response against lung viral infections(*42, 43*). Both decrease in barrier integrity(*39, 40, 44*), and decreased microbicidal activity of myeloid cells (*45–47*) are known to influence superinfection sensitivity. Therefore, we analyzed the relative contribution of IFN-λ to directly modulate immune functions and epithelial cell biology to the establishment of morbidity and *S. aureus* superinfection sensitivity in response to poly (I:C) stimulation. To this end we generated reciprocal bone marrow chimeras in which either the hematopoietic compartment (*Ifnlr1*^−/−^ → WT) or the stromal compartment (WT → *Ifnlr1*^−/−^) were defective for IFN-λ signaling, and the appropriate controls (*Ifnlr1*^−/−^ → *Ifnlr1*^−/−^, WT → WT). WT → *Ifnlr1*^−/−^ mice phenocopied complete *Ifnlr1*^−/−^ → *Ifnlr1*^−/−^ chimeras, displaying lower barrier damage before and after *S. aureus* infection (Fig. 2G, and Fig. S5), and decreased bacterial burden (Fig. 2H). In contrast, *Ifnlr1*^−/−^ → WT behaved like WT → WT mice (Fig. 2G, H). While IFN-λ may modulate immune function indirectly (*71*), we did not observe differences in myeloid immune cell recruitment in *Ifnlr1*^−/−^ mice compared to WT (fig. S6A-D). Moreover, depletion of neutrophils did not impact bacterial burden under our experimental conditions (Fig. S6E), further supporting a direct role of IFN-λ on the epithelial barrier. Overall, these data demonstrate that IFN-λ signaling in epithelial cells is necessary and sufficient to impair lung tolerance, inducing barrier damage and susceptibility to a secondary infection.

In order to determine the molecular mechanisms elicited by IFN-λ in epithelial cells, we performed a targeted transcriptomic analysis on sorted epithelial cells from either WT or *Ifnlr1*^−/−^ mice after poly (I:C) treatment (Fig. 3A, S7). Pathway analysis of differentially expressed genes (DEGs), revealed a potent downregulation of the IFN signature in *Ifnlr1*^−/−^ compared to WT epithelial cells (Fig. 3B), confirming the predominant role of this group of IFNs compared to IFN-I during prolonged viral stimulation in the lung. Consistent with the observed defect in barrier function, genes associated with apoptosis and the activation of the p53 pathway (Fig. 3B) were enriched in WT epithelial cells compared to *Ifnlr1*^−/−^ (Fig. 3B), while pathways involved in positive regulation of the cell cycle were enriched in *Ifnlr1*^−/−^ cells (Fig. 3C). Accordingly, epithelial cells in *Ifnlr1*^−/−^ mice proliferate more efficiently after poly (I:C) administration (Fig. 3D) (as well as after *S.aureus* superinfection (Fig. 3E)) as measured by 5-Ethynyl-2’-deoxyuridine (EdU) incorporation, or expression of the proliferation marker Ki67 (Fig. 3F). In agreement with these data, epithelial cells in WT → *Ifnlr1*^−/−^ chimeras also proliferate more efficiently than mice that bear WT stromal cells, and phenocopy *Ifnlr1*^−/−^ → *Ifnlr1*^−/−^ chimeras (Fig. 3G). These data further support the direct activity of IFN-λ on epithelial cells.

**Figure 3.**
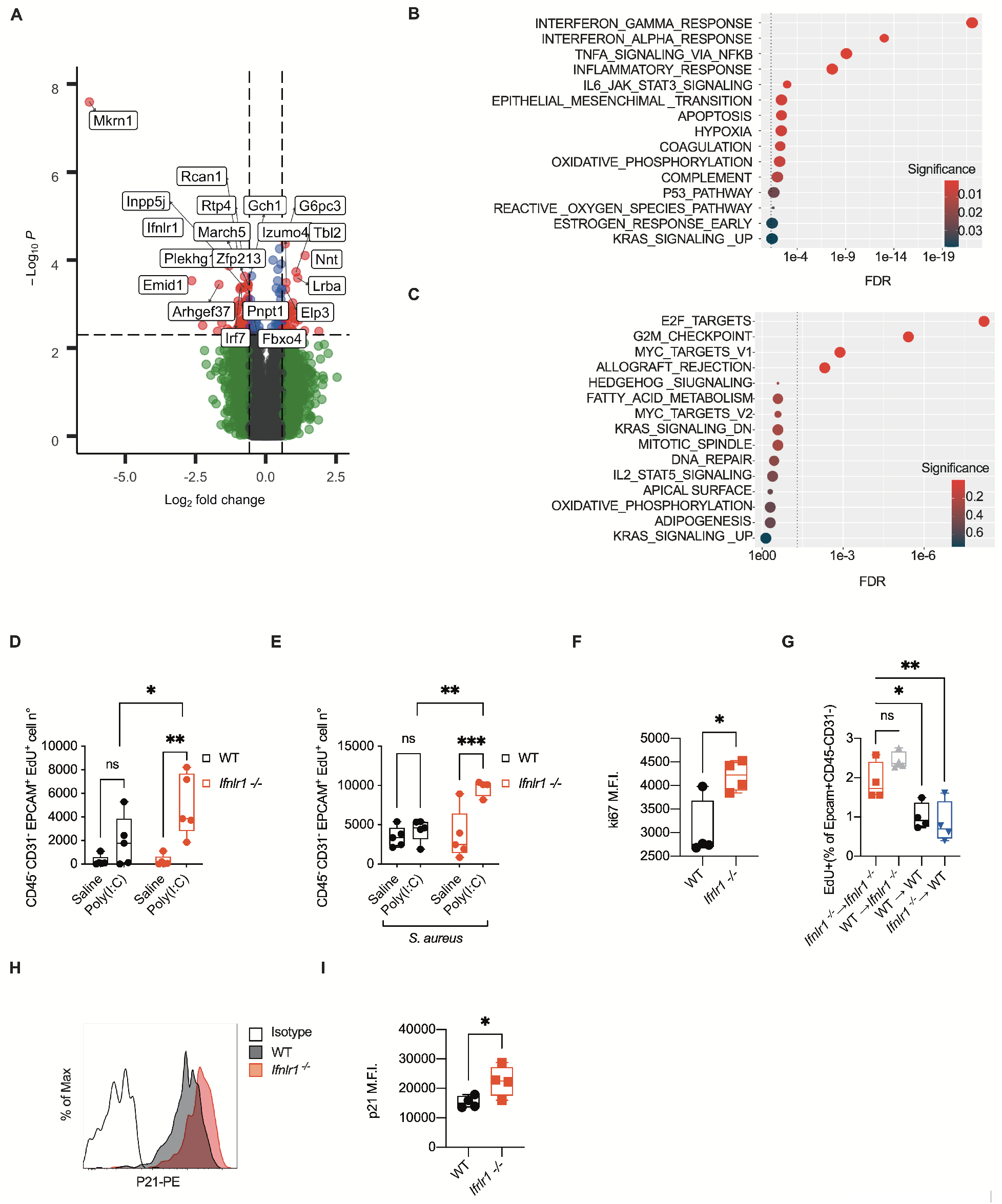
IFN-λ signaling directly inhibits lung epithelia proliferation and impairs repair upon viral recognition. (A-C) Targeted transcriptome sequencing was performed on lung epithelial cells isolated on day 6 from WT and *Ifnlr1*^−/−^ mice treated daily with i.t. 0.5 mg/kg poly (I:C) for 6 days. (A) Volcano plot of differentially expressed genes (DEGs) between WT and *Ifnlr1*^−/−^. DEGs (p-value < 0.005) with a Fold Change > 1.5 (or < −1.5) are indicated in red. Non-significant DEGs (P-value > 0.005) are indicated in green. (B-C) Dot Plot visualization of Gene Set Enrichment Analysis (GSEA) for pathways enriched in (B) WT epithelial cells compared to *Ifnlr1*^−/−^ and (C) *Ifnlr1*^−/−^ epithelial cells compared to WT. The color of the dots represents the adjusted p-value (Significance) for each enriched pathway and its size represents the percentage of genes enriched in the total gene set. (D, E) Epithelial cell proliferation was assessed as EdU incorporation in lung epithelial cells (CD45-CD31-EPCAM^+^) in WT and *Ifnlr1*^−/−^ mice treated as in (A-C) (D) or treated as in (A-C) and infected on day 6 i.t. with 0.5 × 10^8^ CFU *S. aureus* for 12h (E). (F) Mean fluorescent Intensity (MFI) of Ki67 in CD45-CD31-EPCAM^+^ cells of WT and *Ifnlr1*^−/−^ mice treated as in (A-C). (G) EdU incorporation in lung epithelial cells of WT or *Ifnlr1*^−/−^ chimeric mice reconstituted with *Ifnlr1*^−/−^ or WT bone marrow treated as in (E). (H, I) p21 levels in lung epithelial cells (CD45-CD31-EPCAM^+^) from WT and *Ifnlr1*^−/−^ mice treated as in (A-C). Representative histogram (H) and MFI (I) are depicted. (A-C) 4 mice per genotype. (D-I) Each symbol represents one mouse. (D-I) Representative data of 3 independent experiments. Statistics: ns, not significant (P > 0.05); *P < 0.05, **P < 0.01 and ***P < 0.001. (D, E) Two-way ANOVA, (G) One-way ANOVA, (F, I), and two-tailed t test (I) were performed.

Interestingly, the gene that was most downregulated in *Ifnlr1*^−/−^ epithelial cells compared to WT cells is the E3 ubiquitin-protein ligase makorin-1 (*Mrkn1*) (Fig. 3A), which controls p53 and p21 stability by favoring their proteasomal degradation (*48*). Under oxidative stress condition and DNA damage, a hallmark of severe viral infections (*49*), p53 is stabilized by phosphorylation and p21 degradation, mediated by *Mkrn1*, favors apoptosis over DNA repair (*48*). Indeed, *Ifnlr1*^−/−^ epithelial cells, that express lower levels of *Mkrn1*, have elevated levels of p21 as measured by flow cytometry (Fig. 3H, I). These data indicate that the capacity of IFN-λ to reduce tissue tolerance stems from its capacity to inhibit tissue repair by directly influencing epithelial cell proliferation and viability. Also, that p21 degradation via *Mrkn1* upregulation is potently influenced by IFN-λ signaling.

RNA viruses can use several strategies to modulate the immune response to their advantage(*33, 50*), therefore it is crucial to understand the molecular pathways involved in the maintenance of sustained IFN-λ production. Moreover, the difference between mRNA expression and protein levels of interferons suggest that a low abundance cell type with high secretory capacity may be responsible for long term IFN-λ production. We thus investigated the cellular source and molecular pathways that drive IFN-λ production in our model. Early after initial influenza virus infection, IFN-λ is expressed by infected epithelial cells, however, at later time points, DCs from the parenchyma of the lung start to express high levels of the IFN-λ transcript(*5*). We thus hypothesized that lung DCs are the main producers of IFN-λ and are responsible for the secretion of IFN-λ during viral infections. Accordingly, sorted lung resident dendritic cells express high levels of IFN-λ transcript after 5 days of poly (I:C) treatment, in contrast to epithelial cells, alveolar macrophages and monocytes (Fig. 4A), which, instead, express IFN-I and pro-inflammatory cytokines (Fig. S8A, B). Moreover, diphtheria toxin (DT)-mediated depletion of CD11c^+^ cells in CD11c-DT receptor (DTR) mice was sufficient to completely abolish IFN-λ transcript and protein upregulation upon 6 days of poly (I:C) treatment (Fig. 4B, C), while IFN-I production remained unaltered (Fig. S8C, D). Having established that DCs are the main cell type involved in IFN-λ production, we analyzed the relevance of the three main RNA virus recognition pathways in the induction of IFN-λ in dendritic cells (i.e. TLR7-MyD88, RIG-I/MDA5-MAVS, TLR3-TRIF). We thus generated DCs from FMS-like tyrosine kinase 3 ligand (FLT3L)-derived bone marrow cell cultures (FLT3L-DCs). *In vitro* derived FLT3L-DCs comprise plasmacytoid DCs (pDCs), cDC1 and cDC2 like cells (*53*) and are well suited to model tissue DC responses *in vitro*. IFN-λ was induced only when the TLR3-TRIF pathway was activated by administering poly (I:C) in the supernatant, while neither stimulation of the RIG-I/MDA5-MAVS pathway by intracellular delivery of poly (I:C), nor stimulation of TLR7 were able to induce IFN-λ (Fig. 4D). Consistently with the response measured *in vivo*, TLR7 stimulation did not induce IFN production while it induced upregulation of pro-inflammatory cytokines, and intracellular delivery of poly (I:C) induced high levels of IFN-I but not IFN-λ (Fig.4D, Fig. S9A, B). In agreement with the key role of TLR3, IFN-λ production upon extracellular poly (I:C) encounter was abolished by genetic deletion of the signaling adaptor TRIF (encoded by the gene *Ticam1*) but not by deletion of the RIG-I/MDA5 adaptor MAVS (*Mavs*) (Fig. 4D). Conversely, IFN-I production in response to intracellular delivery of poly (I:C) was largely dependent on the signaling adaptor MAVS (Fig. S9A). Consistent with our previous data, when the RIG-I/MAVS pathway was activated by transfection of the influenza A virus derived pathogen-associated molecular pattern (PAMP) 3-phosphate-hairpin-RNA (3p-hpRNA), IFN-I but not IFN-λ, was efficiently induced in a MAVS-dependent manner (Fig. S10A-E, poly (I:C) was used as a control). Finally, inhibition of endosomal acidification by treatment with the pharmacological agent chloroquine abolished IFN-λ induction in response to extracellular poly (I:C), while it preserved IFN-I production upon intracellular poly (I:C) delivery (Fig. S11A, B). These evidences clearly indicate that TLR3 stimulation potently induces IFN-λ production by DCs *in vitro*. We, thus, explored the importance of the TLR3-TRIF pathway *in vivo* under our experimental conditions. Dendritic cells sorted from *Ticam1*^−/−^ mice treated with poly (I:C) for six days did not express appreciable levels of IFN-λ transcripts while still produced type I interferons (Fig. 4E, F). Moreover, poly (I:C) treated *Ticam1*^−/−^ mice were protected from *S. aureus* superinfections (Fig. 4G), and the decrease in bacterial burden correlated with lower IFN-λ transcript levels in the lung, although IFN-I levels remained similar to those of WT mice (Fig. 4H, I). Confirming the crucial role of TLR3 signaling in DCs for IFN-λ production, chimeric mice in which *Ticam1*^−/−^ bone marrow (BM) cells are transferred in a WT irradiated host (*Ticam1*^−/^→WT) phenocopied *Ticam1*^−/−^ animals (Fig. 4J-L).

**Figure 4.**
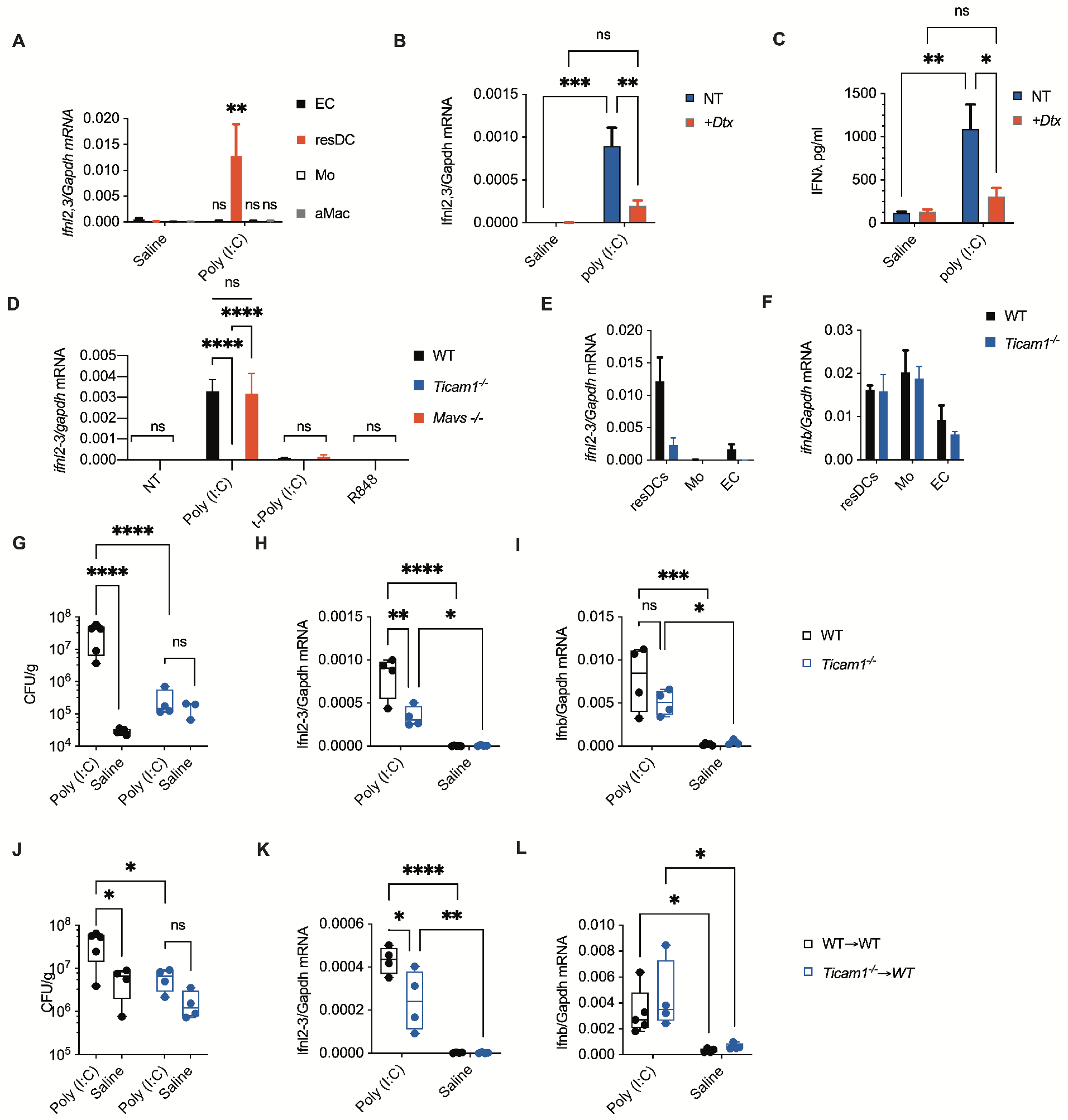
Lung resident DCs produce IFN-λ in a TRIF-dependent manner upon viral recognition. (A) *Ifnl2,3* relative mRNA expression in sorted lung epithelial cells (EC), resident DC (resDC), monocytes and monocyte-derived cells (Mo) and alveolar macrophages (aMac) sorted from WT mice treated daily with i.t. 0.5 mg/kg poly (I:C) or saline for 6 days was measured on day 6. (B) CD11c-DTR mice were injected with diphtheria toxin (DTx) to deplete the CD11c* cells *in vivo*. Relative *Ifnl2,3* mRNA (B) and IFN-λ protein levels (C) from lung homogenates were evaluated on day 6. (D) FLT3-DCs from WT, *Ticam1*^−/−^ or *Mavs*^−/−^ mice were treated with 50μg/ml poly (I:C), 1 μg/ml transfected poly (I:C) or 50μg/ml R848 for 3h. Relative *Ifnl2,3* mRNA expression was evaluated by qPCR. *Ifnl2,3* (E) and *Ifnb1* (F) relative mRNA expression in sorted lung epithelial cells (EC), resident DC (resDC), and monocytes and monocyte derived cells (Mo) sorted from WT and *Ticam1*^−/−^ mice treated as in (A) was measured on day 6. (G-I) WT and *Ticam1*^−/−^ mice were treated with poly (I:C) as in (A) and subsequently infected with i.t. 0.5 × 10^8^ CFU of *S. aureus* on day 6 for 12h. Lung bacterial burden normalized by lung weight (G), *Ifnl2,3* (H) and *Ifnb1* (I) relative mRNA expressions were evaluated. (J-I) WT chimeric mice reconstituted with *Ticam1*^−/−^ bone marrow (*Ticam1*->WT) or WT bone marrow (WT->WT) were treated as in (G-I). Lung bacterial burden normalized by lung weight (J), *Ifnl2,3* (K) and *Ifnb1* (L) relative mRNA expressions 12 hpi were evaluated. Representative data of 3 independent experiments. Statistics: ns, not significant (P > 0.05); *P < 0.05, **P < 0.01 and ***P < 0.001 (Two-way ANOVA). (G-L) Each dot represents one mouse. Median and range are depicted. (A-F) Mean and SEM of 4 mice (A-C, E, F) and of 3 independent experiments (D) are depicted.

The immune system evolved to prevent and resist to pathogen invasion but doing so often threatens host fitness and causes disease in the form of immunopathology (*51*). RNA viruses are the major cause of most severe lower respiratory tract viral infections (*52, 53*). While most virus infections manifest as self-limiting upper respiratory tract infections, influenza viruses, SARS-CoV, SARS-CoV-2 and MERS-CoV can progress to severe lung disease with potentially lethal outcomes(*50, 54, 55*). Although different viruses vary in their virulence and pathogenic potential, the most severe cases of lung RNA viral infections share similar features that suggest an immune pathological etiology. In COVID-19, SARS, MERS and flu, severe symptoms and death occur late after the initial symptoms onset, and after the peak in viral load (*56–61*) further indicating a central role for an immune etiology of the most severe forms.

While IFN-λ is uniquely equipped to induce a gentler immune response that favors viral clearance in the lungs without inducing overt immune activation (*1, 3, 62*), its impact on epithelial cell biology and its effect on the maintenance of tissue integrity and tolerance to pathogen invasion is incompletely understood. In a system that allowed us to isolate the effect of immune activation from resistance to viral infection, we demonstrate that sustained IFN-λ production in the lung in response to viral PAMPs compromises epithelial barrier function, induces lung pathology and morbidity and predisposes to lethal secondary infections by impairing the capacity of the lungs to tolerate bacterial invasion. Loss of lung barrier tolerance is sufficient to induce lethality upon bacterial challenge independently of bacterial growth (*39*), and alteration of the repair response in the lung can favor bacterial invasion independently from immune cell control (*63*). In our model immune cell recruitment is not affected by IFN-λ and neutrophils are dispensable for the impaired control of bacterial infections, while IFN-λ signaling on epithelial cells is necessary and sufficient to cause heightened bacterial invasion.

Under our experimental conditions, TLR3-TRIF signaling in conventional lung DCs is responsible for the induction of IFN-λ. This is consistent with reports indicating that *Tlr3*-deficient mice are protected from influenza-induced immune pathology(*64*). Moreover, TLR3 detects replication intermediates from necrotic cells (*35*) and is, thus, insensitive to viral immune evasion. This is of particular interest during highly pathogenic human coronavirus infections, whose success in establishing the initial infection is partly due to their ability to dampen TLR7 and MAVS dependent early IFN responses(*50*), as indicated by their poor ability to induce IFN responses *in vitro* in epithelial cells (*31*). Indeed, severe clinical manifestations and surge of IFN and cytokine production are evident after the initial infection has peaked, as demonstrated by the analysis of the BAL of ICU-hospitalized COVID-19 patients.

IFN-λ has recently been proposed as a therapeutic agent for COVID-19 infections (*13*), and clinical trials have been initiated for mild and early SARS-CoV-2 infections (*14*) by virtue of its unique antiviral and immune modulatory activity. However, our discovery reveals not only that the most severe COVID-19 patients present high levels of type III IFNs in the lower airways, but also a previously unappreciated capacity of IFN-λ to interfere with repair processes, which renders treatment of severe COVID-19 patients with IFN-λ extremely dangerous. By contrast, existing clinical trials are exploring inhibition of the type II cytokine signaling adaptors Janus Kinases (JAKs) in severe COVID-19 patients (*65*). In light of our discoveries, inhibition of JAK kinases could be successful by both inhibiting detrimental activities of proinflammatory cytokines such as IL-6 and IFN-I, and mitigating the antiproliferative effect of IFN-λ. Intriguingly, *in vitro* treatment of dendritic cells with chloroquine, a controversial therapeutic proposed against COVID-19(*66–68*), completely blocks TLR3-dependent IFN-λ production. Whether this inhibitory effect translates in therapeutic settings, and whether it could have beneficial outcomes in COVID-19, remains an open question.

Collectively we demonstrate that sustained activation of antiviral immunity in the lungs induces the production over time of high levels of IFN-λ by DCs through the TLR3-TRIF pathway. Prolonged IFN-λ stimulation interferes with tissue repair by inhibiting epithelial cell proliferation and favoring p53 mediated apoptosis. This inhibitory effect toward barrier repair induced by IFN-λ can drive immunopathogenesis in severe viral infections and predispose the host to impaired tolerance against opportunistic bacterial infections.

## Acknowledgments

We thank Dr. JC Kagan for discussion, help and support.

## Authors contributions

AB, SG, and BS designed, performed, and analyzed the experiments; AB wrote the paper; RS performed the analysis of the sequencing data; FB and ALC performed the experiments; NC, MDS, and NM performed and analyzed human experiments; FG contributed to the design of the experiments; IZ conceived the project, designed the experiments, supervised the study and wrote the paper.

## Data and materials availability

all data is available in the manuscript or the supplementary materials and/or upon request to the corresponding author.

## Materials and Methods

### Mice

C57BL/6J (Jax 00664) (wild-type; WT), B6.SJL-Ptprca Pepcb/BoyJ (*CD45.1*; Jax 002014), C57BL/6J-*Ticam1^Lps2^*/J (*Ticam1*^−/−^; Jax 005037), *B6;129-Mavs^tm1Zjc^*/J (*Mavs*^−/−^;Jax 008634, *B6.FVB-1700016L21RikTg*(*Itgax-DTR/EGFP*)*57Lan*/J (CD11c-DTR, Jax 004509) mice were purchased from Jackson Labs. C57BL/6 IL-28R^−/−^ (*Ifnlr1*^−/−^) mice were provided by Bristol-Myers Squibb. Mice were housed under specific pathogen-free conditions at Boston Children’s Hospital. *Staphylococcus aureus* infections were conducted in the Biosafety Level-2 facility at Boston Children’s Hospital. All procedures were approved under the Institutional Animal Care and Use Committee (IACUC) and conducted under the supervision of the department of Animal Resources at Children’s Hospital (ARCH).

### Reagents and antibodies

Antibodies used for flow cytometry and sorting experiments: CD45.1 (A20), CD45.2 (104), total CD45 (30-F11), EpCAM (G8.8), CD31 (MEC13.3), CD24 (M1/69), MHC-II I-A/I-E (M5/114.15.2), Ly6G (1A8), Ly6C (HK1.4), CD11b (M1/70), CD11c (N418), CD64 (X54-5/71), Siglec F (S17007L), CD3 (45.2C11), CD19 (6D5), erythroid cell marker (Ter119), Ki67 (16A8), purchased from Biolegend; p21 (F-5) purchased from Santa Cruz. Live/dead cell markers Zombie Red^™^ (423109) or Zombie Violet^™^ (423113) dyes were purchased from Biolegend.

For *in vitro* and/or *in vivo* studies, poly (I:C) HMW (tlr-pic), R848 (tlr-r848) and 3p-hpRNA (tlrl-hprna) were purchased from Invivogen. For *in vivo* administration of type III IFN, we used polyethylene glycol-conjugated IFN-λ2 (PEG-IFN-λ) (gift from Bristol-Myers Squibb). Diphtheria toxin (unnicked) from *Corynebacterium diphtheriae* was purchased from Cayman Chemical. Anti-Ly6G antibody, clone 1A8 (BE0075-1) and rat IgG2a isotype control (BE0089) for *in vivo* administration was purchased from Bioxcell. 2’-Deoxy-5-ethynyl uridine (EdU) was purchased from Carbosynth (NE08701). Chloroquine (PHR1258), Fluorescein isothiocyanate (FITC)-Dextran (MW:10,000da) (FD10S), Deoxyribonuclease (DNase) I from bovine pancreas (DN25) and Dispase II (D4693) were purchased from MilliporeSigma.

### *In vivo* treatments and infection

Intratracheal instillations (i.t.) were performed as previously described in (*69*) 0.5mg/kg of poly (I:C) HMW, R848 or saline were administered i.t. daily for 6 days or as indicated in the figure legends. Where indicated, mice were treated i.t. with recombinant 50μg/kg PEG-IFN-λ concomitantly from day 2 post-treatment, with R848 stimulation. *Staphylococcus aureus* subsp. *aureus* Rosenbach (ATCC^®^ 25904^™^) was grown at 37°C in Tryptic Soy Broth (TSB) for 16h to the autolytic phase, then subcultured and grown to an optical density (OD_600_) of 0.4, centrifuged and resuspended in PBS immediately prior to infection. Each mouse was infected by i.t. instillation of 0.5 × 10^8^ CFU of *S. aureus* and sacrificed 12 hours post-infection, except for in survival studies. Rectal temperature and body weights were monitored daily.

### Survival study and endpoints

Mice were deemed to have reached endpoint at 75% of starting weight or after reaching body temperature of 25°C or lower.

### Generation of bone-marrow chimeras

To generate mice with hematopoietic-specific deletion of *Ifnlr1* or *Ticam1*, 6-week-old CD45.1 + mice were exposed to lethal whole-body irradiation (950 rads per mouse) and were reconstituted with 5 × 10^6^ donor bone marrow cells from 6-week-old wild-type, *Ifnlr1*^−/−^ or *Ticam1*^−/−^ mice. Mice were treated with sulfatrim in the drinking water and kept in autoclaved cages for 2 weeks after reconstitution. After 2 weeks, mice were placed in cages with mixed bedding from wild-type, and *Ifnlr1*^−/−^ or *Ticam1*^−/−^ mice to replenish the microbiome and were allowed to reconstitute for 2 more weeks. A similar procedure was used to generate bone-marrow chimeras with stromal cells-specific deletion in *Ifnlr1*. Here, recipient WT or *Ifnlr1*^−/−^ mice underwent irradiation and were reconstituted with BM cells derived from CD45.1+ mice similarly as described above.

To evaluate the percentage of chimerism, a sample of peripheral blood was taken from chimeric mice after 4 weeks of reconstitution and stained for CD45.1 and CD45.2 (antibodies as identified under ‘Reagents and antibodies’) and were analyzed by flow cytometry.

### Depletion of Dendritic cells and Neutrophils

In order to deplete CD11c^+^ cells, CD11c-DTR mice received 16μg/kg diphtheria toxin (DTx) intravenously starting one day before TLR ligand or saline administration and continuing every other day until day 6 post-treatment to maintain depletion.

*In vivo* depletion of neutrophils was carried out by injecting anti-Ly6G antibody (100μg/mouse) intraperitoneally, starting one day before treatments and then continuing every other day through the duration of the treatment. As controls for no depletion, mice were injected with rat IgG isotype control.

### Barrier permeability assessment

To assess lung permeability, treated mice were administered FITC-dextran (10μg/mouse) i.t. before or after *S. aureus* infection. After 1hr of dextran instillation, blood was collected from the retro-orbital sinus, and the plasma was separated by centrifugation. Leakage of dextran in the bloodstream was measured as FITC fluorescence in the plasma compared to plasma from mock-treated mice.

### Bronchoalveolar lavage (BAL) and lung collection

BAL was collected as described in (*70*) Briefly, the lungs of euthanized mice were lavaged through the trachea with 3ml PBS to collect the BAL. Samples were centrifuged and the supernatants were used for total protein measurement (Pierce BCA Protein Assay, Thermo Fisher Scientific #23227) and LDH quantification (Pierce LDH Cytotoxicity Assay, Thermo Fisher Scientific #C20301). Lungs were excised and used for RNA extraction using TRI Reagent (Zymo Research #R2050-1-200).

### Bacterial load and lung cytokine production measurement

The left lobe of the lung was weighed and homogenized in 1ml of sterile D.I. water in a Fisherbrand^™^ Bead Mill 24 Homogenizer. To calculate bacterial load, homogenate was serially diluted and plated on TSB-Agar plates in duplicate. Colonies were counted after 16h incubation, and bacterial burden in the lungs was calculated as CFU normalized to individual lung weight. Cytokines production in the lungs was measured in the supernatants collected after centrifuging the lung homogenates.

### Flow cytometry and cell sorting

Lung cells were isolated as described in (*71*) Briefly, mice were euthanized and perfused. 2 ml of warm dispase solution (5mg/ml) were instilled into the lungs followed by 0.5ml of 1% low-melt agarose (Sigma #A9414) at 40°C, and allowed to solidify on ice. Inflated lungs were incubated in dispase solution, for 30’ at RT. The lungs were then physically dissociated, incubated 10’ with DNAse I 50 μg/ml and filtered through 100μm and 70μm strainers. Red blood cells were lysed using ACK buffer. Single cell suspensions were stained for live/dead using Zombie Red or Zombie Violet, and then with antibodies against surface antigens diluted in PBS + BSA 0.2% for 20 minutes at 4°C. Cells were then washed, fixed with 3.7% paraformaldehyde for 10 minutes at room temperature, washed again and resuspended in PBS + BSA 0.2%. Samples were acquired on a BD LSRFortessa flow cytometer and data were analyzed using FlowJo v.10 software (BD Biosciences). CountBright Absolute Counting Beads (Invitrogen #C36950) were used to quantify absolute cell numbers.

For cell sorting, FACS samples were prepared as described above, and sorted on Sony MA900 Cell Sorter following the sorting strategy indicated in Fig. S12. The sorted cells were collected directly into TRI Reagent for RNA extraction. For detecting cell proliferation, fixed cells were treated to assess for 5-Ethynyl-2’-deoxyuridine (EdU) incorporation as described below (‘Epithelial cell proliferation’) before being stained with antibodies against cell-surface antigens. Intracellular staining of Ki67 and p21 were carried out using FoxP3 Fix/Perm Buffer set (Biolegend #421403) following the manufacturer’s instructions.

### Epithelial cell proliferation

Proliferation of lung epithelial cells was monitored by assessing the incorporation of EdU (100 mg/kg, intraperitoneally, 12h before end-point euthanasia), and analyzed by flow cytometry or histology. Briefly, single cell suspensions of lung cells from mice injected with EdU were isolated as described before (‘Flow cytometry and cell sorting’). Single cell suspensions were assessed for live/dead and fixed using 3.7% PFA. Staining for EdU was carried out by Click-chemistry reaction. Fixed cells were permeabilized with 0.1% Triton X100 (Millipore-Sigma) for 15 min. After permeabilization cells were washed and incubated with 4 mM Copper sulphate (Millipore-Sigma), 100 mM Sodium ascorbate (Millipore-Sigma) and 5 μM sulfo-Cyanine3-azide (Lumiprobe #A1330) in Tris Buffered Saline (TBS) 100mM, pH 7.6, for 30 min at room temperature.

### Ion Torrent

For targeted transcriptome sequencing, 20 ng of RNA isolated from sorted cells was retro-transcribed to cDNA using SuperScript VILO cDNA Synthesis Kit (ThermoFisher Scientific). Barcoded libraries were prepared using the Ion AmpliSeq Transcriptome Mouse Gene Expression Kit as per the manufacturer’s protocol and sequenced using an Ion S5 system (ThermoFisher Scientific). Differential gene expression analysis was performed using the Transcriptome Analysis Console (TAC) software with the ampliSeqRNA plugin (ThermoFisher Scientific).

Genes were called expressed (n=11,294) if they had average log2 expression of 2 or greater in either WT or *Ifnlr1*^−/−^. Differentially expressed genes (DEGs) between WT and *Ifnlr1*^−/−^ were selected by thresholding on fold change (+/-1.5) and *p* value (0.005). In heatmaps, DEGs were Z-scaled and clustered (Euclidean distance, Ward linkage). Pathway analysis was performed with the R package hypeR, using Gene Set Enrichment Analysis on genes ranked according to their Log2(Fold Change).

### Cell culture

FLT3L-DCs were differentiated from bone marrow cells in Iscove’s Modified Dulbecco’s Media (IMDM; Thermo Fisher Scientific), supplemented with 30% B16-FLT3L derived supernatant and 10% fetal bovine serum (FBS) for 9 days.

Differentiated cells were stimulated with poly (I:C), R848, or 3p-hpRNA. Where indicated poly (I:C) and 3p-hpRNA were transfected with Lipofectamine 300 (Invitrogen #L3000015) according to manufacturer’s instructions at the concentrations indicated in the figure legends. Where indicated poly (I:C) stimulated cells were pre-treated with 10μg/ml chloroquine for 5 minutes prior to stimulations.

### qRT-PCR and ELISA

RNA was isolated from cell cultures using a GeneJET RNA Purification Kit (Thermo Fisher Scientific #K0731) according to manufacturer’s instructions. RNA was extracted from excised lungs by homogenizing them in 1ml of TRI Reagent. RNA was isolated from TRI Reagent samples using phenol-chloroform extraction or column-based extraction systems (Direct-zol RNA Microprep and Miniprep, Zymo Research #R2061 and #R2051) according to the manufacturer’s protocol. RNA concentration and purity (260/280 and 260/230 ratios) were measured by NanoDrop (ThermoFisher Scientific).

Purified RNA was analyzed for gene expression on a CFX384 real-time cycler (Bio-Rad) using a TaqMan RNA-to-CT 1-Step Kit (Thermo Fisher Scientific) or SYBR Green (Bio-Rad). Probes specific for *Ifnl2/3, Ifnb1, Il1b, Rsad2, Gapdh* were purchased from Thermo Fisher Scientific, and SYBR-Green primers for *Rsad2, Cxcl10, Gapdh* were purchased from Sigma. Cytokine analyses were carried out using homogenized lung supernatants, and cell supernatants from stimulated FLT3L-DCs. IFNλ2/3 ELISA (R&D Systems DY1789B) and mouse IFNβ, IL1β, IL-6, TNFα ELISA (Invitrogen) were performed according to manufacturer’s instructions.

### Clinical samples

Bronchoalveolar lavages (BAL) were obtained from five intensive care unit (ICU)-hospitalized SARS-CoV-2-positive patients. In parallel, five naso-oropharyngeal swabs were collected from both SARS-CoV-2-positive and -negative subjects. Among positive patients, two were hospitalized but without the need of ICU support, whereas three out of five did not require any hospitalization. The negative swabs were obtained from subjects undergoing screening for suspected social contacts with COVID-19 subjects. Swabs were performed by using FLOQSwabs^®^ (COPAN) in UTM^®^ Universal Transport Medium (COPAN). All samples were stored at −80°C until processing. The study involving human participants was reviewed and approved by San Raffaele Hospital IRB in the COVID-19 Biobanking project. The patients provided written informed consent.

### RNA extraction protocol and Real-Time PCR of clinical samples

RNA extraction was performed by using PureLink^™^ RNA Thermo Fisher Scientific according to manufacturers’ instruction. In particular, 500 μL for each BAL and swab analyzed sample were lysed and homogenized in the presence of RNase inhibitors. Then ethanol was added to homogenized samples which were further processed through a PureLink^™^ Micro Kit Column for RNA binding. After washing, purified total RNA was eluted in 28 μL of RNase-Free Water. Reverse transcription was then performed according to SuperScript^™^ III First-Strand Synthesis System (Invitrogen^™^) protocol by using 8 μL of RNA extracted from each BAL and swab sample. qRT-PCR analysis for was then carried out for evaluating IL6, IL1B, IFNB1, IFNA2, IFNL1 and IFNL2 expression. All transcripts were tested in triplicate for each sample by using specific primers. GAPDH was also included. Real-time analysis was then performed according to manufacturer instructions by using TaqMan^®^ Fast Advanced Master Mix (Applied Biosystems^™^ by Thermo Fisher Scientific). Real-Time PCR Analysis was performed on ABI 7900 by Applied Biosystems.

### Statistical Analyses

Statistical significance for experiments with more than two groups was tested with one-way ANOVA, and Dunnett’s multiple-comparison tests were performed. Two-way ANOVA with Tukey’s multiple-comparison test was used to analyze kinetic experiments. Two-way ANOVA with Sidak’s multiple-comparison test was used to analyze experiments with 2 grouped variables (i.e. treatment, genotype). Statistical significance for survival curves were evaluated with the Log-rank (Mantel-Cox) test and corrected for multiple comparisons with Bonferroni’s correction. To establish the appropriate test, normal distribution and variance similarity were assessed with the D’Agostino-Pearson omnibus normality test using Prism8 (Graphpad) software. When comparisons between only two groups were made, an unpaired two-tailed *t*-test was used to assess statistical significance. To determine the sample size, calculations were conducted in nQuery Advisor Version 7.0. Primary outcomes for each proposed experiment were selected for the sample size calculation and sample sizes adequate to detect differences with an 80% power were selected. For animal experiments, four to ten mice per group were used, as indicated in the figure legends.

---

**Figure S1.**
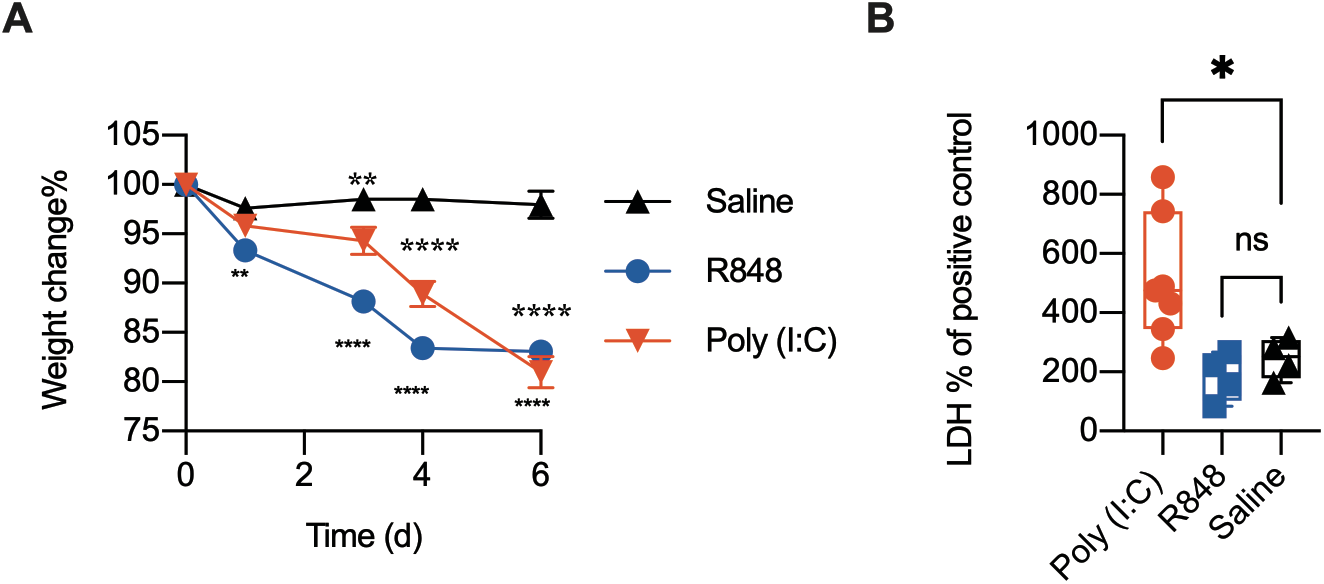
Intratracheal poly (I:C) treatment induces morbidity and lung damage. (A) Weight change and (B) LDH released in BAL of WT mice treated daily with i.t. 0.5 mg/kg poly (I:C), 0.5 mg/kg R848 or saline for 6 days. Statistics: ns, not significant (P > 0.05); *P < 0.05, **P < 0.01 and ***P < 0.001 (A) Two-way ANOVA, (B) One-way ANOVA. (A) Mean and SEM of 5 mice per group is depicted. (B) Each mouse represents one point. Median and range are depicted.

**Figure S2.**
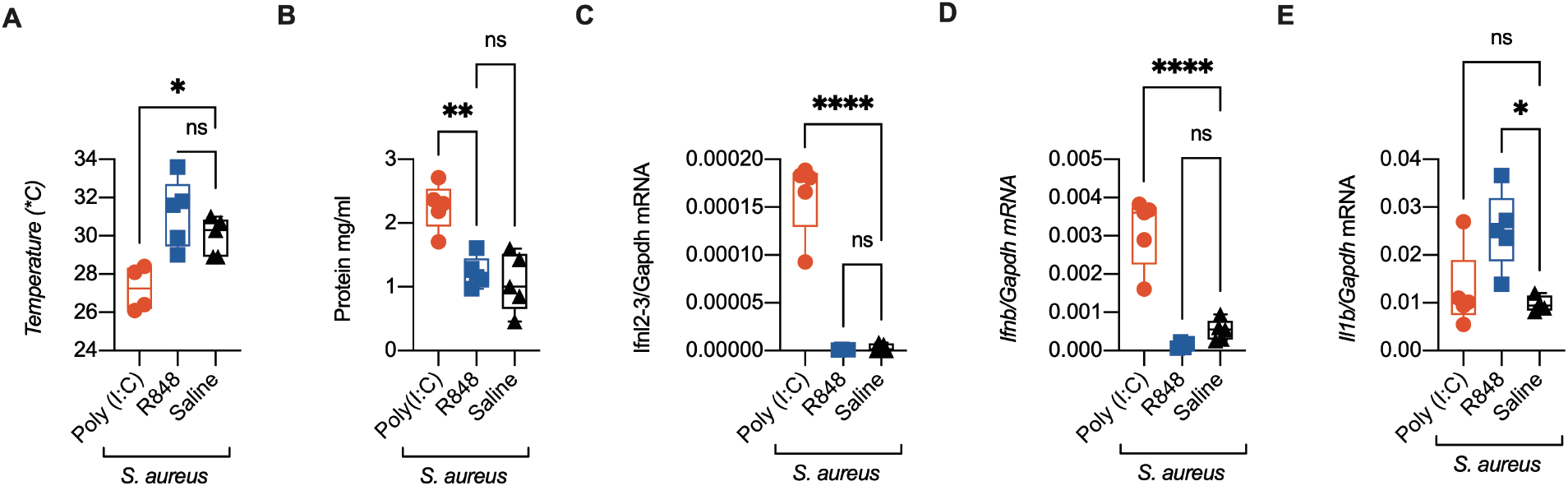
Intratracheal poly (I:C) treatment increases lung susceptibility to bacterial infection. WT mice were treated daily with i.t. 0.5 mg/kg poly (I:C), 0.5 mg/kg R848 or saline for 6 days and infected i.t. with 0.5 × 10^8^ CFU of *S. aureus* at day 6. (A) Body temperature, (B) total protein in the BAL and *Ifnl2,3* (C), *Ifnb1* (D), *Il1b* (E) relative mRNA expression in total lung lysates were evaluated 12 hpi. Statistics: ns, not significant (P > 0.05); *P < 0.05, **P < 0.01 and ***P < 0.001 (One-way ANOVA). Each mouse represents one point. Median and range are depicted.

**Figure S3.**
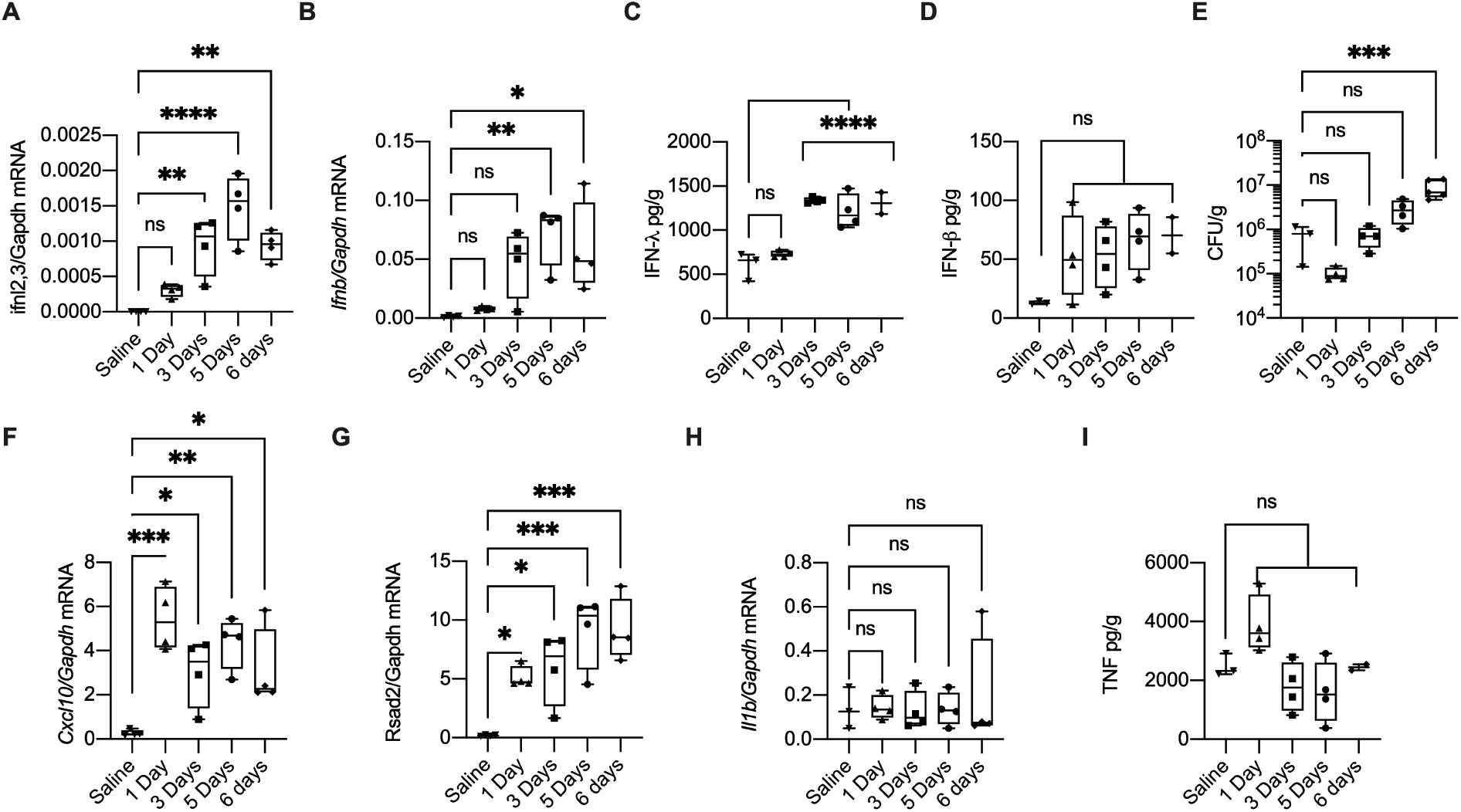
IFN-λ protein levels correlate with susceptibility to bacterial infections. (A-H) WT mice were treated daily for 1, 3, 5 or 6 days with i.t. 0.5 mg/kg poly (I:C) or 6 days of saline, and infected with i.t. 0.5 × 10^8^ CFU of *S. aureus* for 12h. Total lung homogenates were analyzed by qPCR for *Ifnl2,3* (A), *Ifnb1* (B), *Cxcl10* (F), *Rsad2* (G), *Il1b* (H) relative mRNA expression. Protein levels of IFN-λ (C), IFN-β (D) and TNF-α (I) were evaluated by ELISA on lung homogenates. (E) Bacterial burden was evaluated in total lung homogenate. Statistics: ns, not significant (P > 0.05); *P < 0.05, **P < 0.01 and ***P < 0.001 (One-way ANOVA compared to Saline treatment). Each mouse represents one point. Median and range are depicted.

**Figure S4.**
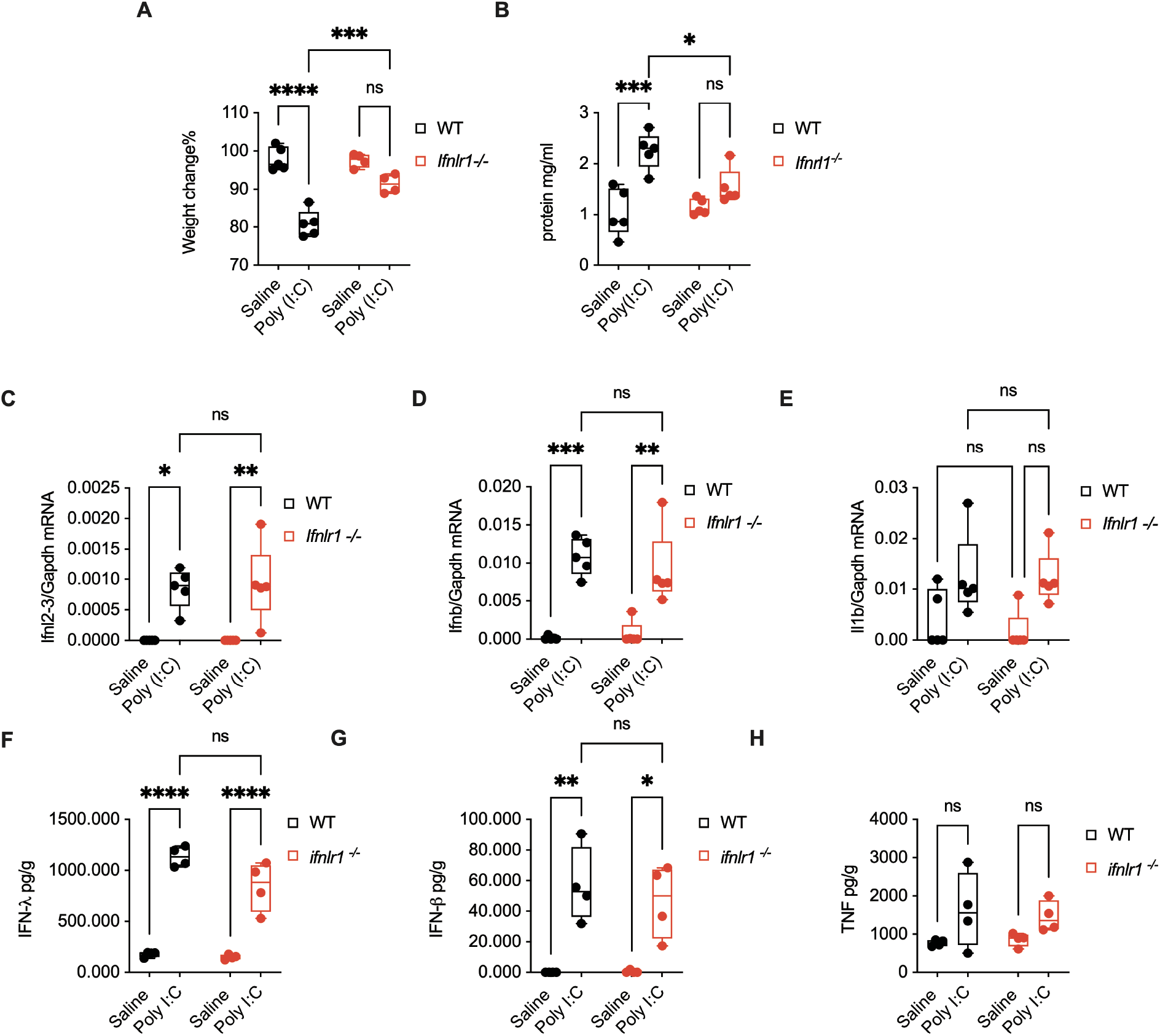
IFN-λ signaling is necessary to confer poly (I:C)-induced morbidity and susceptibility to bacterial infections. WT and *Ifnlr1*^−/−^ mice were treated daily with i.t. 0.5 mg/kg poly (I:C) for 6 days and infected with i.t. 0.5 × 10^8^ CFU of *S. aureus* for 12h. (A) Weight change, (B) total protein in the BAL, *Ifnl2,3* (C), *Ifnb1* (D), *Il1b* (E) relative mRNA expression, and IFN-λ (F), IFN-β (G) and TNF (H) production in total lung homogenate were evaluated. Statistics: ns, not significant (P > 0.05); *P < 0.05, **P < 0.01 and ***P < 0.001 (Two-way ANOVA). Each mouse represents one point. Median and range are depicted.

**Figure S5.**
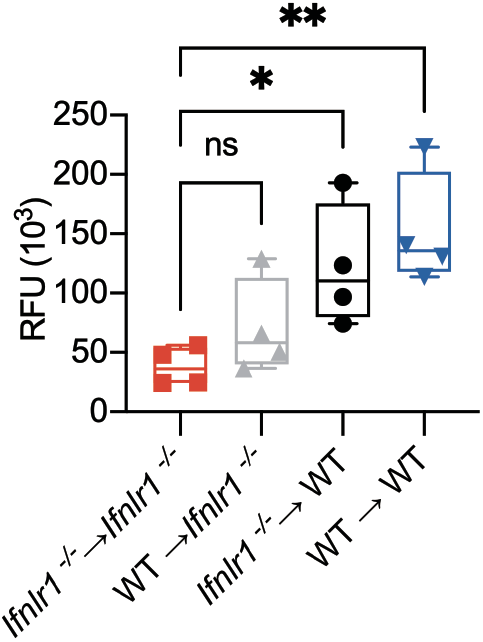
IFN-λ acts on epithelial cell to induce lung damage. Barrier function of chimeric mice *Ifnlr1*^−/−^→*Ifnlr1*^−/−^, WT→*Ifnlr1*^−/−^, *Ifnlr1*^−/−^→WT, and WT→WT were measured as RFU of FITC-dextran in the plasma of mice that were treated daily with i.t. 0.5 mg/kg poly (I:C) for 6 days and infected with i.t. 0.5 × 10^8^ CFU of *S. aureus* for 12h. Statistics: ns, not significant (P > 0.05); *P < 0.05, **P < 0.01 and ***P < 0.001 (One-way ANOVA). Each mouse represents one point. Median and range are depicted.

**Figure S6.**
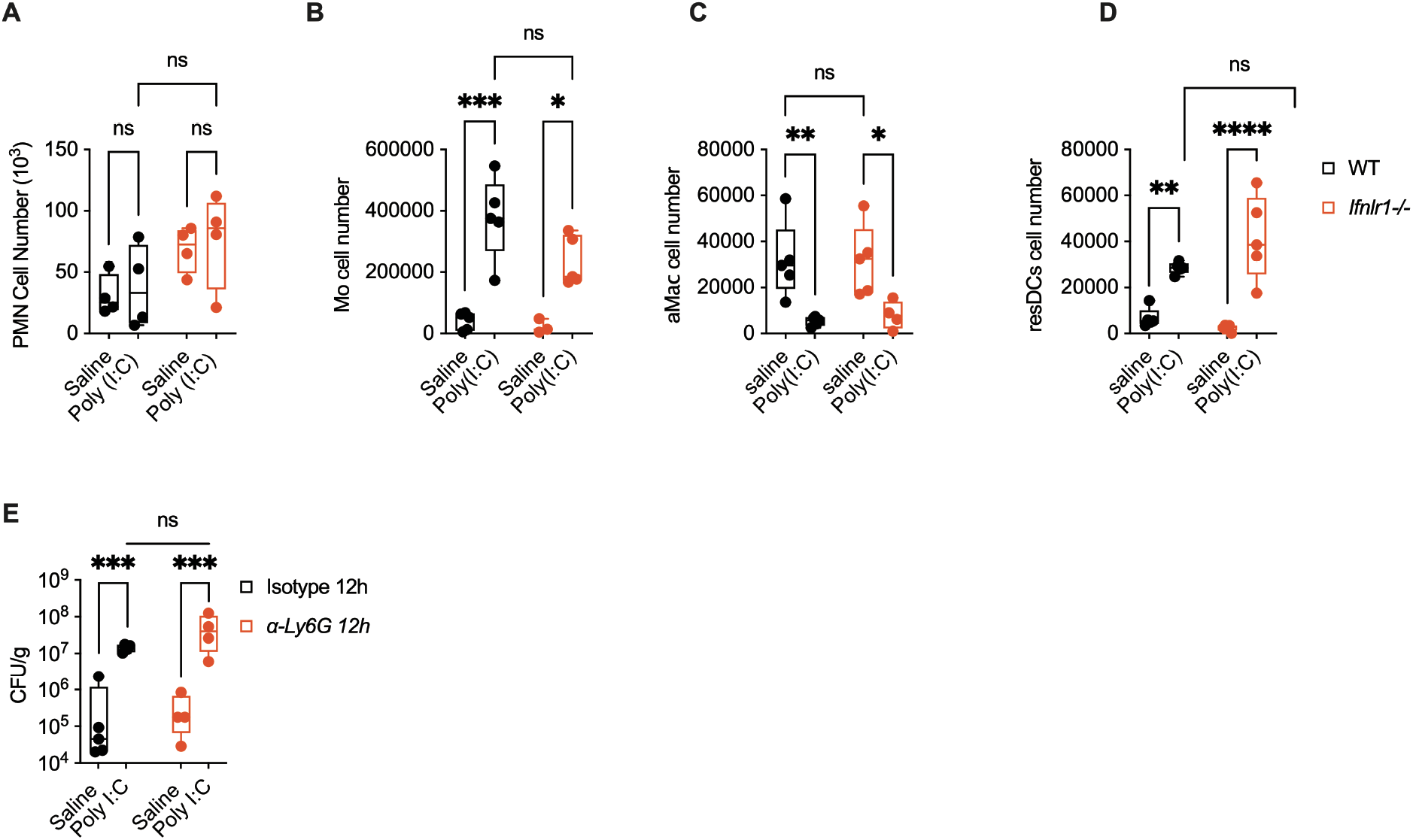
IFN-λ dependent loss of tolerance against bacterial infections is independent of neutrophil function or alteration in immune cell recruitment. Total number of (A) neutrophils (PMN), (B) Monocyte and monocyte derived cells (Mo), (C) alveolar macrophages (aMac) and (D) resident DC (resDCs) in lungs of WT and *Ifnlr1*^−/−^ mice treated daily with i.t. 0.5 mg/kg poly (I:C) or saline for 6 days and infected with *S. aureus* as in Fig. 1K. (E) Wild-type mice were treated with either isotype or anti-Ly6G antibody to deplete Ly6G^+^ neutrophils *in vivo*. Mice were treated with poly (I:C) or saline for 6 days and infected with i.t. 0.5 × 10^8^ CFU of *S. aureus* for 12h. Bacterial loads in the lungs were calculated as CFU per gram of lung weights. Statistics: ns, not significant (P > 0.05); *P < 0.05, **P < 0.01 and ***P < 0.001 (Two-way ANOVA). Each mouse represents one point. Median and range are depicted. Logarithmic data are fitted when depicting Bacterial load (E)

**Figure S7.**
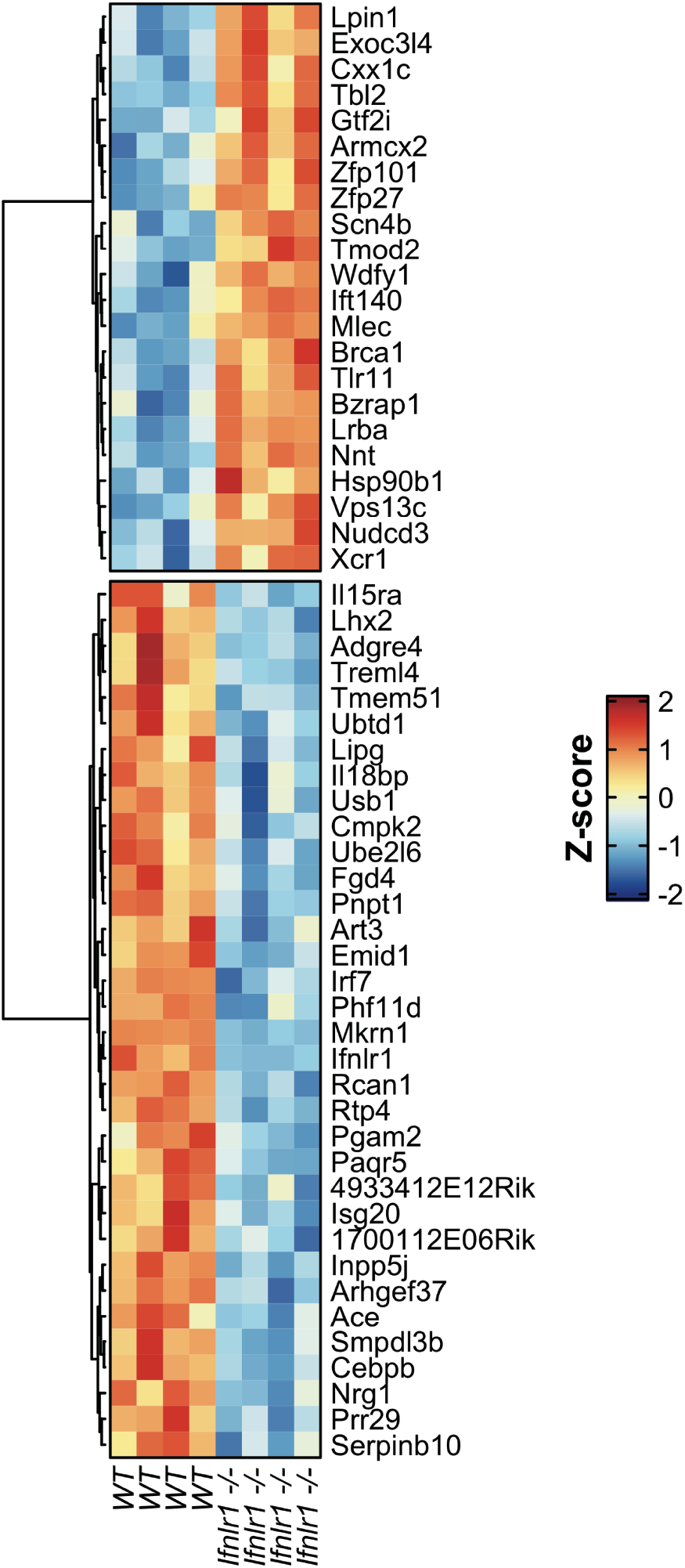
Differentially expressed genes in WT versus *Ifnlr1*^−/−^ lung epithelial cells from poly (I:C)-treated mice. Heatmap of genes with a p-value < 0.005 and a Fold Change greater than 1.75 (or lower than −1.75) between *Ifnlr1*^−/−^ and WT lung epithelial cells.

**Figure S8.**
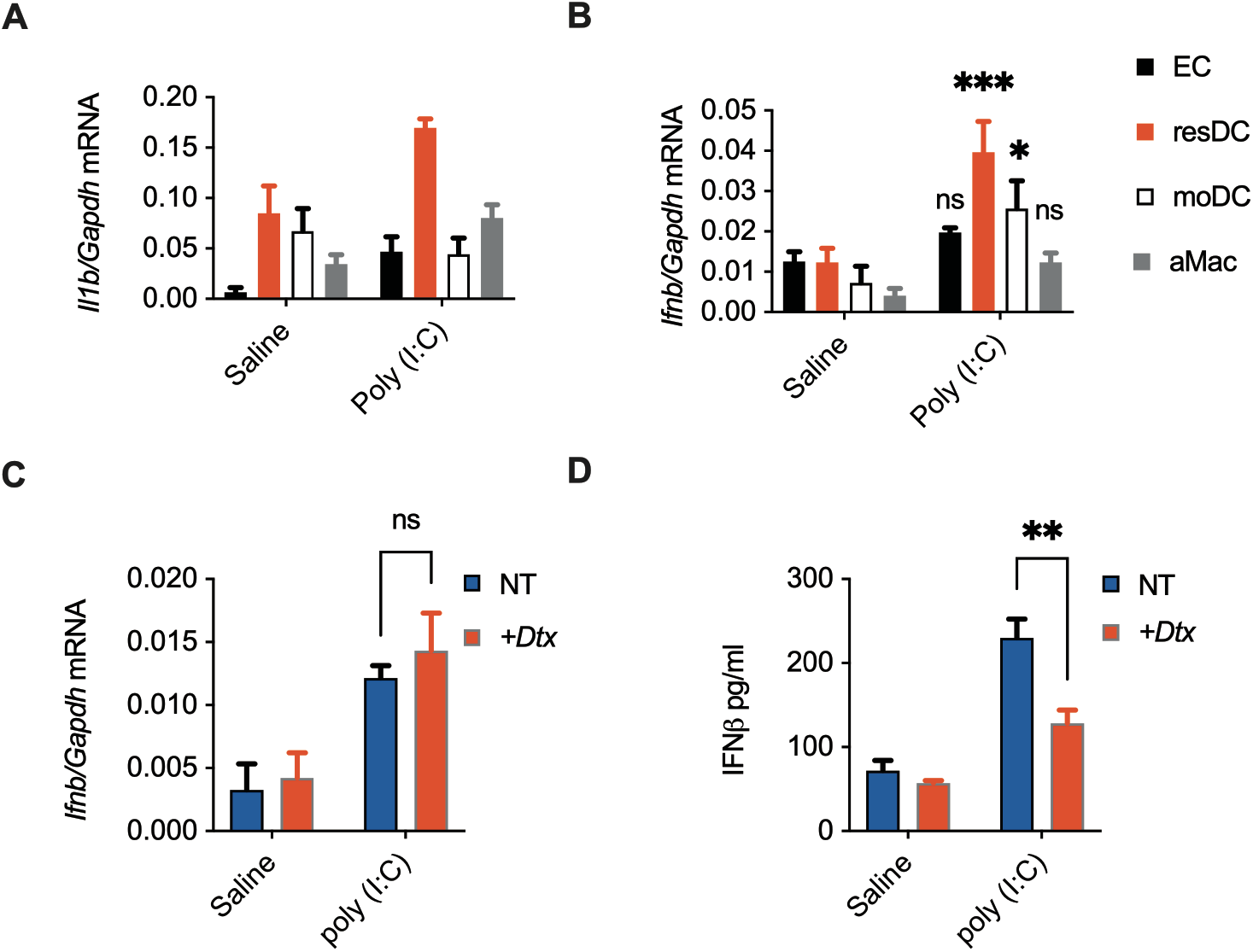
Lung resident DCs are the primary producers of IFN-β upon poly (I:C) treatment. Lungs of mice treated daily with i.t. 0.5 mg/kg poly (I:C) or saline for 6 days were sorted for epithelial cells (EC), resident DC (resDC), monocyte-derived DC (moDC), and alveolar macrophages (aMac) and assessed for (A) *Il1b* and (B) *Ifnb1* relative mRNA expressions. CD11c-DTR mice depleted for CD11 c^+^ cells *in vivo* by DTx injections were treated daily with i.t. 0.5 mg/kg poly (I:C) or saline for 6 days. Total lung lysates of the treated mice were analyzed for (C) *Ifnb1* relative mRNA expression, and (D) IFN-β protein expression by ELISA. Statistics: ns, not significant (P > 0.05); *P < 0.05, **P < 0.01 and ***P < 0.001 (Two-way ANOVA). Mean and SEM of 5 mice per group are depicted.

**Figure S9.**
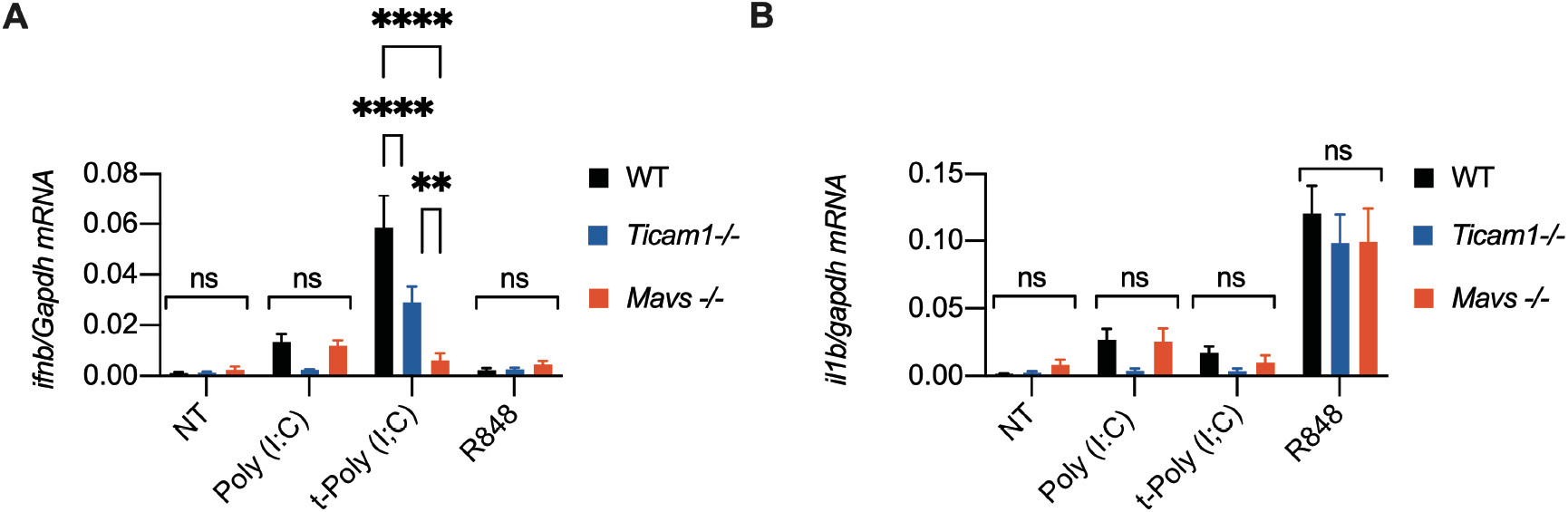
FLT3L-DCs responses to TLR3, RIG-I or TLR7 ligands. FLT3L-DCs from WT, *Ticam1*^−/−^ or *Mavs*^−/−^ mice were treated with 50μg/ml poly (I:C), 1μg/ml transfected poly (I:C) or 50μg/ml R848 for 3h. *Ifnb1* (A), and *Il1b* (B) relative mRNA expressions were evaluated by qPCR. Statistics: ns, not significant (P > 0.05); *P < 0.05, **P < 0.01 and ***P < 0.001 (Two-way ANOVA). Mean and SEM of 3 independent experiments is depicted.

**Figure S10.**
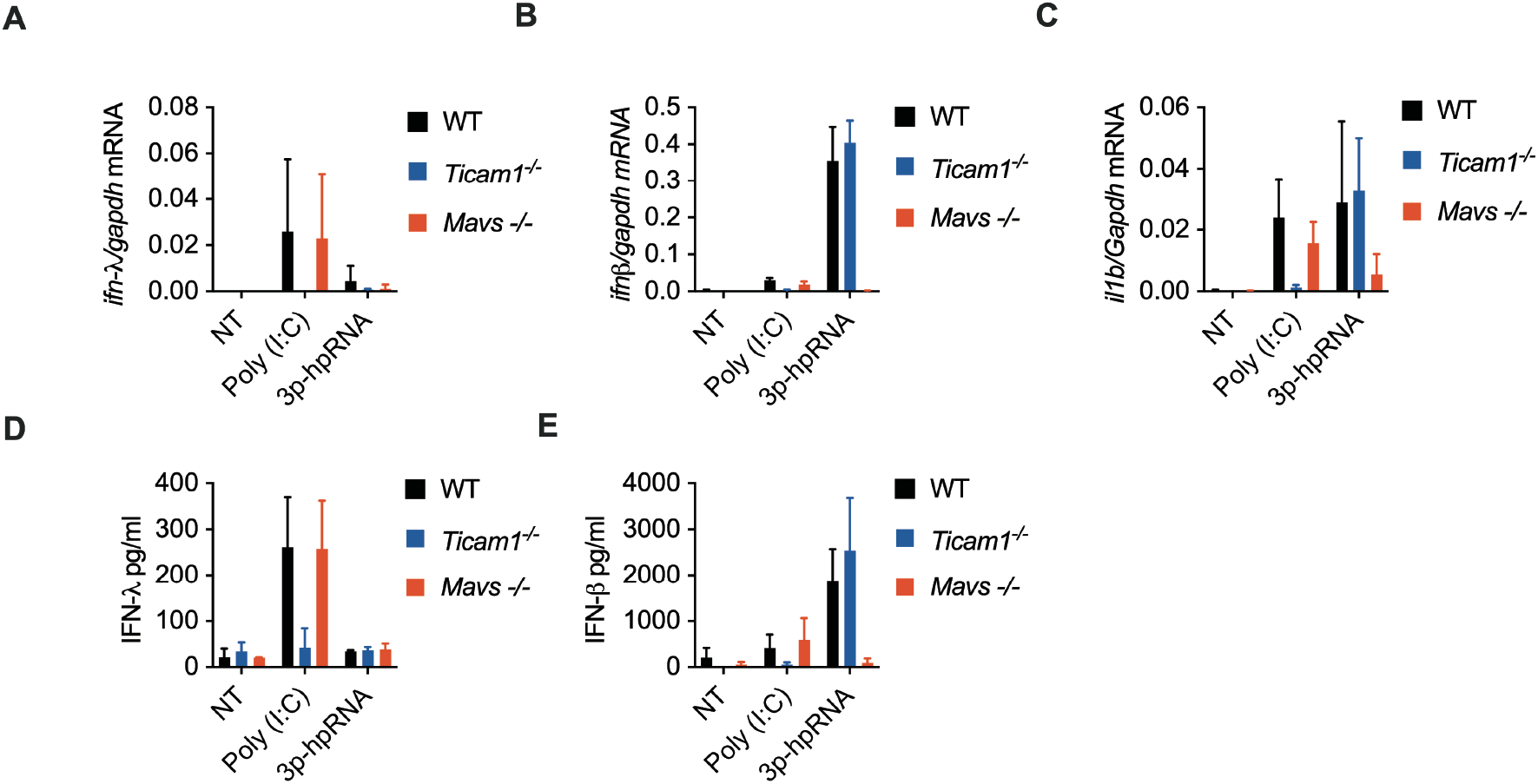
FLT3L-DCs upregulate IFN-λ uniquely upon activation of TLR3 signaling and not in response to the RIG-I specific ligand 3p-hpRNA. FLT3L-DCs from WT, *Ticam1*^−/−^ or *Mavs*^−/−^ mice were treated with 50μg/ml poly (I:C), or 1μg/ml transfected 3p-hpRNA for 3h or 6h. *Ifnl2,3* (A), *Ifnb1* (B), and *Il1b* (C) relative mRNA expressions were evaluated by qPCR after 3h. IFN-λ (D), and IFN-β (E) levels in the supernatants were evaluated by ELISA after 6h.

**Figure S11.**
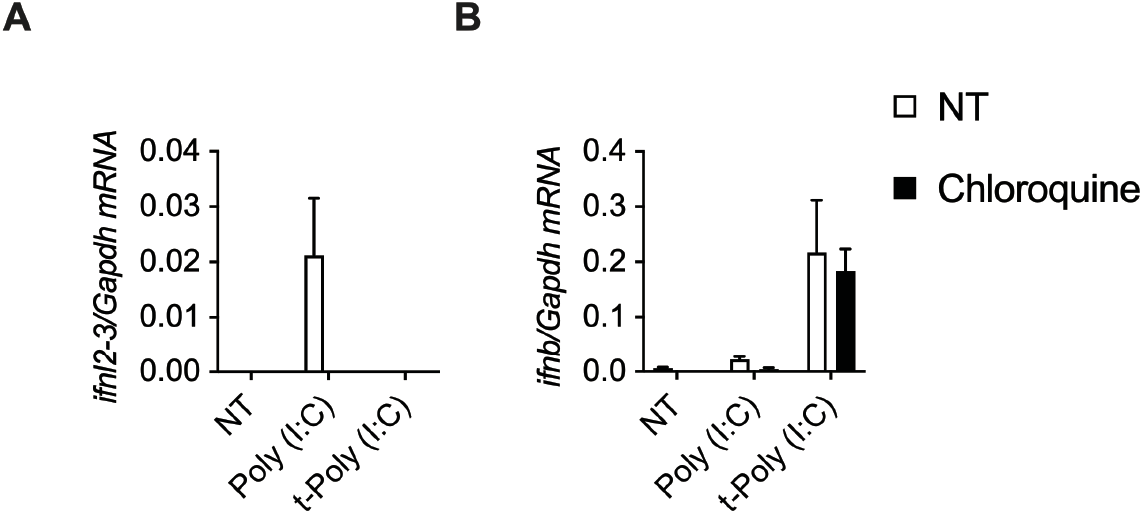
The endosomal TLR inhibitor Chloroquine inhibits poly (I:C) dependent IFN-λ expression in FLT3L-DCs. FLT3L-DCs from WT mice were treated with 50μg/ml poly (I:C), or 1μg/ml transfected poly (I:C) for 3h in the presence or absence of 10μg/ml Chloroquine. *Ifnl2,3* (A), and *Ifnb1* (B) relative mRNA expressions were evaluated by qPCR.

**Figure S12.**
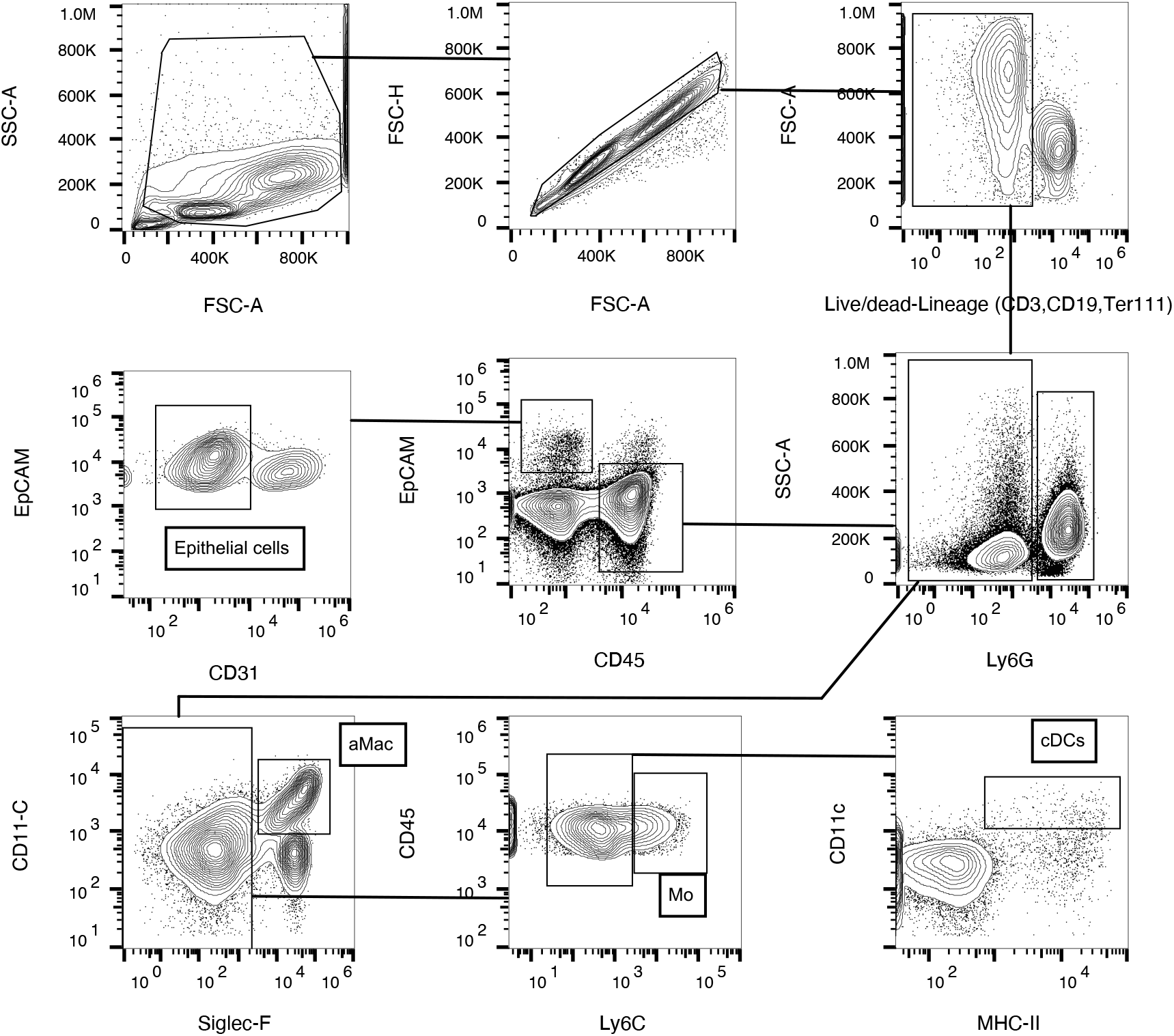
Sorting and cytofluorimetry strategy. Depiction of gating strategy for cell sorting. Cells were gated on FSC and SSC to eliminate debris, on FSC-A -FSC-H to select single cells and cells negative for live/dead dye and Lineage markers (CD3, CD19, Ter119). Epithelial cells were gated as CD45^−^ EpCAM^+^CD31^−^. The EpCAM^−^ cells were sorted for immune cells as follows: aMac were gated as CD45^+^Ly6g^−^CD11c^hi^Siglec-F^+^, monocytes and monocyte-derived cells (Mo) were gated as CD45^+^Ly6g^−^Siglec-F^−^Ly6C^+^, cDCs were gated as CD45^+^Ly6g^−^Siglec-F^−^ Ly6C^−^CD11c^+^MHC-II^hi^.

